# Enhanced Viral Metagenomics with Lazypipe 2

**DOI:** 10.1101/2022.12.21.521450

**Authors:** Ilya Plyusnin, Olli Vapalahti, Tarja Sironen, Ravi Kant, Teemu Smura

## Abstract

Viruses are the main agents causing emerging and re-emerging infectious diseases. It is therefore important to screen for and detect them and uncover the evolutionary processes that support their ability to jump species boundaries and establish themselves in new hosts. Metagenomic next-generation sequencing (mNGS) is a high-throughput, impartial technology that has enabled virologists to detect either known or novel, divergent viruses from clinical, animal, wildlife and environmental samples, with little a priori assumptions. mNGS is heavily dependent on bioinformatic analysis, with an emerging demand for integrated bioinformatic workflows. Here, we present Lazypipe2, an updated mNGS pipeline with, as compared to Lazypipe1, significant improvements in code stability, transparency, with added functionality and support for new software components. We also present extensive benchmarking results, including evaluation on a novel canine simulated metagenome, and precision and recall of virus detection at varying sequencing depth and low to extremely low proportion of viral genetic material. Additionally we report accuracy of virus detection with two strategies: homology searches using nucleotide or amino acid sequences. We show that Lazypipe2 with nucleotide-based annotation approaches near perfect detection of eukaryotic viruses and, in terms of accuracy, outperforms the compared pipelines. We also discuss the importance of homology searches with amino acid sequences for the detection of highly divergent novel viruses.

## 1. INTRODUCTION

Fast development in metagenomic Next-generation sequencing (mNGS) and analysis has enabled virologists to assess the true diversity of viruses in clinical, animal, wildlife and environmental samples. mNGS is a high-throughput, impartial technology with many advantages compared to established diagnostic methods for virus detection [1]. mNGS can detect viruses that do not propagate in cell cultures and, unlike the PCR- or antigen-based detection, can detect a broad spectrum of viruses without a priori assumptions on the likely targets. Overall, mNGS has a potential to translate into universal methods for virus discovery, surveillance and broad-spectrum clinical diagnostics [1–3]. That said, we should note that mNGS is a relatively novel technology and there are still challenges to address such as accessibility, costs and sampling to reporting time.

There is a growing interest within virology to utilise mNGS, specifically in the detection of viruses that cannot be cultured [4,5]. There is also a growing interest for applications in clinical settings, particularly for difficult to diagnose cases with rare or unknown disease aetiologies that would otherwise require multiple targeted tests [6,7]. mNGS is also recognised for its potential for the monitoring and early detection of emerging viral pathogens [1].

mNGS approaches are heavily dependent on bioinformatic analysis that processes raw sequence output by the NGS sequencer into metagenomic assemblies and various reports on the microorganisms nucleic acid presence and abundances in the analysed samples. Generally, analysis of NGS data requires bioinformatic expertise, computational resources and, in many cases, installation and maintenance of large reference databases [2,3,7]. This has raised concerns that the lack of such in the public health laboratories or smaller research facilities can cause hurdles for the adoption of mNGS methods [2,3,7]. These challenges can be addressed by developing bioinformatic pipelines and services designed to handle bioinformatic and resource-related challenges in mNGS sequence analysis. During the last decade many pipelines for virus discovery and sample composition analysis have emerged: VMGAP [8], PathSeq [9], VIROME [10], READSCAN [11], VirusFinder [12], SURPI [13], MetaVir [14], VIP [15], MetaShot [16], VirusSeeker [17], viGEN [18], Genome Detective [19], Kraken2 [20] and IDseq [2]. mNGS pipelines tested in clinical settings are also starting to emerge [7].

Our research group has contributed to this development with the Lazypipe mNGS pipeline designed primarily for virus discovery from clinical, animal and environmental samples with minimal requirements in terms of bioinformatic expertise and/or resources [21]. Lazypipe has been adopted as a standard module for mNGS analysis at the Finnish IT Center for Science (www.csc.fi) and has been successfully applied to detect and characterise a multitude of novel viral pathogens from a variety of different sample types, including arthropod vectors [22,23], mosquitos [24], farm animals [25] and wildlife [26]. Here, we present an enhanced version of our mNGS pipeline, Lazypipe2. In this latest version of our pipeline we introduce code updates to achieve better installation experience, stability, speed and transparency as well as a smaller memory and disk-space footprint. We also added new functionalities to support new analysis options and further automation of frequent user cases. New features include support for SPAdes assembler [27], minimap2 annotations [28], 2nd round annotation verification with NCBI blastn [29], support for massively parallel execution for large sample batches and support for snakemake interface [30]. Furthermore, we compiled a novel canine simulated metagenome and performed an extensive benchmarking on human and canine simulated metagenomes. Also, using our benchmarks we analysed errors in virus detection and were able to detail sources of these errors, as well as, compare nucleotide and amino acid based annotation strategies.

## 2. MATERIAL AND METHODS

### 2.1 Code updates

#### 2.1.1 Code restructuring for better stability and transparency

To improve the portability of Lazypipe1 (all versions freely available at https://bitbucket.org/plyusnin/lazypipe/), we excluded several external Perl modules that posed installation challenges from our users. Among other things, we removed dependencies on spreadsheet modules and BioPerl modules. Spreadsheet modules were replaced with the R openxlsx library. All sequence manipulations with BioPerl were reimplemented with SeqKit calls [31]. All parsing and handling of taxonomic paths were reimplemented with TaxonKit calls [32]. To improve stability and transparency, all manipulations with tab-separated value (tsv) files were reimplemented using column headers instead of indices. To work with named headers we used csvtk toolkit [33]. All three tools mentioned here (SeqKit, TaxonKit and csvtk toolkit) were selected based on similar criteria, namely simple installation without dependencies, fast binary implementation and support for multithreading. These and other updates are illustrated in Figure 1.

**Figure 1.**
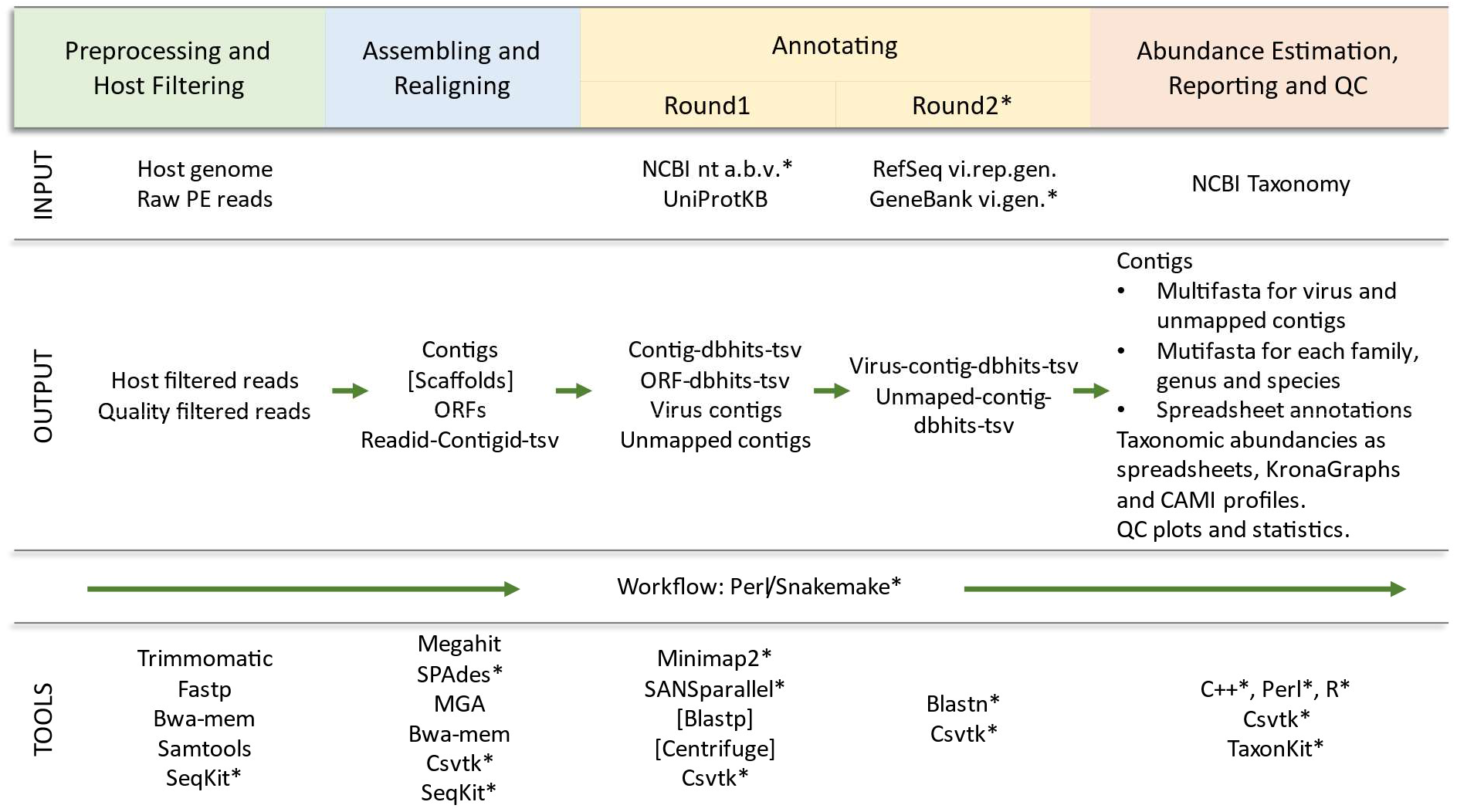
Overview of Lazypipe2 workflow and updates. *Tools and databases that were added or updated in this release compared to the previously published work [21]. INPUT, input files and reference database; dbhits, database homologs returned by the search; NCBI nt abv, viral, bacterial and archaeal entries from the NCBI nt database; RefSeq vi.rep.gen., RefSeq representative virus genomes; GeneBank vi.gen., GeneBank complete virus genomes.

Integration of Lazypipe with SANSparallel was improved by adding a taxonomy operator to the distributed Pannzer package (version 3.0) [34]. This operator handles mapping of UniProtKB accession ids to NCBI taxon ids on the SANSparallel server, which removed the requirement to do this mapping locally by loading into memory large accession to taxon id dictionaries. This reduced the size of the code and also reduced required memory.

In the new version we added printing of collective multi-fasta files for contigs mapped to viruses and contigs with no mapping (*contigs_vi.fa* and *contigs_un.fa*, respectively). The pipeline will also print collective multi-fasta files for each family, genus, and species found in the data.

#### 2.1.2. Support for parallel analysis of large data collections

We also added the generation of unique working directories for each invocation of the pipeline, which added support for parallel analysis of multiple input fastq files. This feature adds support for massive parallel execution of our pipeline, for example with Slurm job arrays. To address the analysis of large data-collections, all output fastq files are now compressed. We also added support for automated cleaning of intermediated files (activated with --*clean option*). These later additions have significantly reduced the disk footprint of the pipeline.

### 2.2 New features

#### 2.2.1. Integrating SPAdes

We added support for read assembling with SPAdes [27], which was shown in several comparative studies to have high performance for simulated and mock community viral metagenomes [14,35].

#### 2.2.2. Integrating minimap2 aligner

Lazypipe1 (version 1.0) supported annotation with both nucleotide (nt) and amino acid (aa) based search engines, using SANSparallel/blastp and Centrifuge, respectively. For the new version, we added support for the minimap2 nt search engine [28]. One practical consideration here was the constantly growing size of the NCBI nt database. For the June 2022 update, our attempt to run the Centrifuge indexer on the bacteria and virus portion of the nt database failed to reach completion after running for 70 hours on 32 cores. Minimap2 was an attractive alternative, since this search supports splitting the database into batches that are indexed “on the fly” as the search progresses. Minimap2 supports assembly to reference alignments (--*x asm5/asm10/asm20* modes) with different expected sequence divergence. By default, Lazypipe2 uses --*xasm20* (approximately 5% divergence), and as our reference we use a custom database covering all viral, bacterial and archaeal entries from the NCBI nt database.

#### 2.2.3. Integrating blastn for 2nd round annotations

The end users will often want to confirm discovered viral genomes using blastn. To support this, we added the --*pipe blastv* option, which will run blastn on contigs identified as virus contigs in the main annotation round. As the reference database, the user can choose any custom or public blastn database. We recommend using RefSeq representative genomes for viruses updated and published by NCBI [29]. We also offer support for a broader virus database that we compiled and updated from GeneBank complete virus genomes (https://bitbucket.org/plyusnin/lazypipe/). We also added support for re-annotation of contigs, which had no database hits in the main annotation round (*contigs_un.fa*). Unmapped contigs can be re-annotated with blastn against a custom database with --*pipe blastu* option.

#### 2.2.4. Improved bacteriophage labelling

We added a more complete labelling of bacterial and archaeal viruses. The new labelling now includes all viral families and orders, including exclusively viral species known to infect bacteria or archaea according to the latest Virus Metadata Resource published by the International Committee on Taxonomy of Viruses (https://ictv.global/taxonomy/ VMR_20-190822). The labelling was made updatable and transparent by listing these bacteriophage families and orders in a separate source file (*R/NGS.phage.filter.R*).

#### 2.2.5. New interface with Snakemake

To support a wider range of users, including those unfamiliar with perl, we complemented the default perl interface with a Snakemake interface [30]. Snakemake is a workflow manager that is able to handle large bioinformatic workflows with complex input-output interdependencies [30].

### 2.3 Benchmarking

We evaluated Lazypipe2 on two benchmarks. For the first benchmark we used the human simulated metagenome from the MetaShot project [16]. For the second benchmark we compiled a novel canine simulated metagenome (described below), based on viruses and bacteria associated with the domestic dog. The domestic dog was chosen in order to test virus detection against a different host and bacterial background. The resulting simulated metagenome can also serve as a valuable tool for future mNGS benchmarking, particularly in the context of companion animal and one-health research.

All benchmarking was done on Linux/Unix CPU supercluster with 32 cores each running at 2.1 GHz.

#### 2.3.1. Human simulated metagenome

Human simulated metagenome was used to evaluate Lazypipe2 (version 2.1) against Lazypipe1 (version 1.0), Kraken2 [20] and CZID [2]. Lazypipe2 was run with default options and minimap2 and SANS aligners. Lazypipe1 was run with default options and SANS aligner. Kraken2 was run with default settings and the standard database. CZID was run via web interface v7.1 with host set to “human” and background to “none”.

#### 2.3.2. Viral genomes associated with the domestic dog

We started by searching RefSeq (version 214) for virus assemblies with host field matching to “*Canis lupus*”, “*Canis lupus familiaris*” or “dog”, or virus name matching “Canine”. This resulted in 44 accessions including 30 complete genomes and 14 complete cds sequences. We then complemented this list by searching VirusHostDB [36] for viruses that were labelled with “*Canis lupus familiaris*” as their host. From this list we manually selected assemblies for viruses that are either well established canine pathogens (e.g. *Lyssavirus rabies*) or that have been isolated from a dog. We further extended our collection by adding *Canine Influenza A virus H3N2* from NCBI Influenza Virus Sequence Database [37]. The H3N2 subtype is the latest and most common Influenza virus isolated from dogs in Asia and the United States [38]. Lastly, we pruned the list of collected canine papillomaviruses to include only one genome for each species-level taxon. This resulted in 7 canine papillomaviruses. The resulting collection of canine viral genomes included 57 assemblies and 39 unique virus taxa (accessions available in File S1).

#### 2.3.3. Bacterial genomes associated with the domestic dog

We also created a collection of bacterial genomes that are associated with the domestic dog. We started by datamining biosample entries from NCBI BioSample database that were labelled with *host_taxid* equal to *9615* or with *host* matching “(*dog[s]?)|(canis lupus familiaris)*”. This returned 21,083 unique dog-associated samples. We then selected from RefSeq genomes database (version 214) all bacterial accessions that were sequenced from dog-associated samples. We further pruned this list to include only unique taxon ids. This resulted in a collection of 195 accessions and 159 bacterial species. We also created a smaller collection that included only complete genomes from the above set (58 accession). Accessions and other details for both canine bacterial collections are available in File S1.

#### 2.3.4. Canine simulated metagenome

To create the canine simulated metagenome we processed our viral, bacterial and host (GCF_000002285.5) sequences with ART [39] uplying settings for Illumina PE 2X 150 nt libraries with HiSeq 2500 built-in profile. Host genome and bacterial genomes were processed with 5X coverage, while viral genomes were processed with coverage ranging from 1X to 5X. We then combined host, bacterial and viral libraries into a number of canine metagenomes with varying proportions of viral and bacterial sequences and varying viral coverage (Table 1). Low proportion/coverage of virus genomes aimed to test virus calling at low to very low abundance of virus genetic material. Canine simulated metagenomes were then used to benchmark taxa calling and read taxonomic binning with Lazypipe2 – *ann minimap*.

**Table 1.** Canine simulated metagenomes. We combined the host library, two bacterial libraries and viral libraries at different coverage levels (5X to 1X).

| Metagenome | Host reads | Bacterial reads | Virus reads | Bacterial taxids | Virus taxids |
| --- | --- | --- | --- | --- | --- |
| dog5X-ba.comp5X-vi5X | 93.37% | 6.60% | 0.021% | 58 | 39 |
| dog5X-ba5X-vi5X | 79.85% | 20.13% | 0.018% | 195 | 39 |
| dog5X-ba5X-vi4X | 79.85% | 20.13% | 0.014% | 195 | 39 |
| dog5X-ba5X-vi3X | 79.86% | 20.13% | 0.011% | 195 | 39 |
| dog5X-ba5X-vi2X | 79.86% | 20.13% | 0.007% | 195 | 39 |
| dog5X-ba5X-vi1X | 79.86% | 20.14% | 0.004% | 195 | 39 |

## 3. RESULTS

### 3.1 Benchmarking taxonomic profiling

#### 3.1.1 Results for human simulated metagenome

We compared Lazypipe2 with minimap2 and SANS aligners to Lazypipe1 with SANS aligner, Kraken2 and CZID on the human simulated metagenome. Precision, recall and F1-score (harmonic mean of precision and recall) for predicted virus and bacterial taxa are given in Tables 2 and 3, respectively. For virus taxa we excluded predictions falling into the last percentile of read distribution and for bacterial taxa into the last 5 percentiles of the read distribution. As discussed previously [21], these settings aim to reduce noise for taxa at lower abundances. For bacterial predictions we also included Lazypipe2 results with the last 20 percentiles excluded (*Lazypipe2 --ann sans -t20*) (Table X.b).

**Table 2.** Accessing accuracy of virus taxon retrieval by different tools. Compared tools are ordered by the descending F1-score. True, number of virus taxa in the benchmark, TP, number of true positive predictions, FP, number of false positive predictions, FN, number of false negative predictions, Pr, precision, Rc, recall, F1, F1-score.

| Tool | Rank | True | TP | FP | FN | Pr | Rc | F1 |
| --- | --- | --- | --- | --- | --- | --- | --- | --- |
| Lazypipe2 --ann minimap | species | 84 | 79 | 2 | 5 | 97.5% | 94.0% | 95.8% |
| Lazypipe2 --ann sans | species | 84 | 75 | 13 | 9 | 85.2% | 89.3% | 87.2% |
| Lazypipe1 --ann sans | species | 84 | 72 | 14 | 12 | 83.7% | 85.7% | 84.7% |
| IDseq | species | 84 | 69 | 12 | 15 | 85.2% | 82.1% | 83.6% |
| Kraken2 | species | 84 | 60 | 19 | 24 | 75.9% | 71.4% | 73.6% |
| Lazypipe2 --ann minimap | genus | 49 | 47 | 0 | 2 | 100.0% | 95.9% | 97.9% |
| IDseq | genus | 49 | 46 | 0 | 3 | 100.0% | 93.9% | 96.8% |
| Lazypipe2 --ann sans | genus | 49 | 44 | 1 | 5 | 97.8% | 89.8% | 93.6% |
| Lazypipe1 --ann sans | genus | 49 | 42 | 1 | 7 | 97.7% | 85.7% | 91.3% |
| Kraken2 | genus | 49 | 35 | 4 | 14 | 89.7% | 71.4% | 79.5% |

**Table 3.** Accessing accuracy of bacterial taxon retrieval by different tools. Compared tools are ordered by the descending F1-score. Here, IDseq predictions were filtered according to the manual (NT_RPM>=10 and NR_RPM>=1). True, number of bacterial taxa in the benchmark, TP, number of true positive predictions, FP, number of false positive predictions, FN, number of false negative predictions, Pr, precision, Rc, recall, F1, F1-score.

| Tool | Rank | True | TP | FP | FN | Pr | Rc | F1 |
| --- | --- | --- | --- | --- | --- | --- | --- | --- |
| Lazypipe2 --ann sans -t20 | species | 71 | 70 | 52 | 1 | 57.4% | 98.6% | 72.5% |
| IDseq | species | 71 | 42 | 24 | 29 | 63.6% | 59.2% | 61.3% |
| Lazypipe2 --ann minimap | species | 71 | 46 | 39 | 25 | 54.1% | 64.8% | 59.0% |
| Lazypipe1 --ann sans | species | 71 | 68 | 128 | 3 | 34.7% | 95.8% | 50.9% |
| Lazypipe2 --ann sans | species | 71 | 70 | 171 | 1 | 29.0% | 98.6% | 44.9% |
| Kraken2 | species | 71 | 37 | 3903 | 34 | 0.9% | 52.1% | 1.8% |
| Lazypipe1 --ann sans | genus | 44 | 44 | 9 | 0 | 83.0% | 100.0% | 90.7% |
| Lazypipe2 --ann sans -t20 | genus | 44 | 44 | 14 | 0 | 75.9% | 100.0% | 86.3% |
| Lazypipe2 --ann minimap | genus | 44 | 35 | 5 | 9 | 87.5% | 79.5% | 83.3% |
| IDseq | genus | 44 | 33 | 3 | 11 | 91.7% | 75.0% | 82.5% |
| Lazypipe2 --ann sans | genus | 44 | 44 | 19 | 0 | 69.8% | 100.0% | 82.2% |
| Kraken2 | genus | 44 | 38 | 1250 | 6 | 3.0% | 86.4% | 5.7% |

Lazypipe2 with minimap2 aligner demonstrated the best accuracy for virus calling with recall at 94.0% and precision at 97.5% (species-level, Table 2). Lazypipe2 with SANS aligner was second best with recall at 89.3% and precision at 85.2%. Both Lazypipe2 variants outperformed Lazypipe1, IDseq and Kraken2 (Table 2).

Lazypipe2 with SANS (*Lazypipe2 --ann sans -t20*) had the overall best accuracy for bacterial calling (98.6% recall and 57.4% precision). Notably, the high precision was achieved by pruning the last 20 percentiles from the result list. This had a large effect by significantly decreasing the false positive bacterial predictions compared to the default pruning of 5 percentiles (*Lazypipe --ann sans* with 29.0% precision). IDseq was second best (59.2% recall and 63.6% precision) followed by *Lazypipe2 --ann minimap*, *Lazypipe1 --ann sans* and Kraken2.

#### 3.1.2. Classification errors for human simulated metagenome

To gain a better understanding of errors in virus detection, we examined in more details the false positive and false negative predictions. We focused on erroneous calls for eukaryotic viruses reported by Lazypipe2 with SANS and minimap2 aligners (Table 4). These represent annotations by sequence homology of nucleotide (nt) contig sequences (minimap2) and amino acid (aa) sequences in open reading frames (SANS). We also refer to these as the nt- and the aa-based annotations, respectively.

**Table 4.** Errors in virus detection from the human simulated metagenome. The table summarises falsepositive (FP) and false-negative (FN) errors (highlighted) reported by Lazypipe2 with *sans* and *minimap2* annotation options. Error causes, represented by the last column, are explained in the text.

| Virus | Lazypipe<br>--ann sans | Lazypipe<br>--ann minimap2 | Error cause |
| --- | --- | --- | --- |
| *Human endogenous retrovirus* | FN | FN | Reads filtered as host reads |
| *Human endogenous retrovirus W* | FN | FN |  |
| *Naples phlebovirus* | FN | FN | Contig (4187 nt) with high (100%) identity to *Toscana phlebovirus* |
| *Mopeia Lassa virus reassortant 29* | FN | FN | Contig (1551 nt) with high (99.9%) identity to *Lassa virus* |
| *Hepatitis B virus* | FN | TP | ORF identification |
| *Uukuniemi uukuvirus* | FN | TP | ORF identification |
| *Influenza A virus* | FN | TP | ORF identification |
| *Mason-Pfizer monkey virus* | FN | TP | Low read count |
| *Toscana phlebovirus* | FP | FP | Contig (4187 nt) with high (100%) identity to FP |
| *Lassa virus* | FP | TN | ORF (1551 nt) with high (100%) identity to FP |
| *Cowpox virus* | FP | TN | ORFs with high (100%) identity to FP |
| *Borna disease virus* | FP | TN | ORFs (429nt, 606 nt and 1113 nt) with high (100%) identity to FP |
| *Central chimpanzee simian foamy virus* | FP | TN | ORFs (668 nt and 696 nt) with high (96.0-99.0%) identity to FP |
| *Eastern chimpanzee simian foamy virus* | FP | TN | ORFs (180-759 nt) with high (98.0-100%) identity to FP |
| *African green monkey simian foamy virus* | FP | TN | ORF (753 nt) with high (81.1%) identity to FP |
| *Human bocavirus* | FP | TN | ORFs (441-2016 nt) with high (100%) identity to FP |
| *Chimeric Tick-borne encephalitis virus/*<br>*Dengue virus 4* | FP | TN | ORF (6108 nt) with high (100%) identity to FP |
| *Phlebovirus SDYY104/China/2011 virus* | FP | TN | ORF (6258 nt) with high (100%) identity to FP |
| *SFTS phlebovirus* | FP | TN | ORF (738 nt) with high (100%) identity to FP |
| *Bocaparvovirus sp.* | FP | TN | ORF (1920 nt) with high (100%) identity to FP |
| *Coronavirus BtRs-BetaCoV/YN2018D* | FP | TN | ORF (966 nt) with high (100%) identity to FP |
| *Uncultured human fecal virus* | TN | FP | 12 contigs (330-586 nt) with high identity to FP |

Erroneous calls were due to a limited number of typical causes (Table 4). Detailed descriptions of all misclassification errors are available in File S2. Here we report the key points.

For retroviruses, false-negatives were caused by host-genome filtering which, in most cases, also removed the retroviral reads. These errors can be avoided by turning host-filtering off.

For the aa-based annotation common false positive errors were due to homologs in the aa space (labelled as “ORF with high identity to FP” in Table 4) and common false negative errors were due to the failure to identify sufficiently long orfs (labelled as “ORF identification” in Table 4). Naturally, these errors did not occur in the nt-based annotations.

There were also cases of false positive homologs having similar or even higher sequence identity to the assembled contigs than the correct genome (labelled as “Contig with high identity to FP”). The latter case may be attributed to random errors introduced by the ART simulation.

#### 3.1.3. Results for canine simulated metagenome

We benchmarked our pipeline on canine simulated metagenome with default options and minimap2 annotation. Pipeline demonstrated 100% recall of virus species for all variants of the canine metagenome except the lowest 1X coverage version for which there was a single false negative (Table 5). Pipeline called a single false positive virus prediction for all metagenome versions. For the dog5X-ba5X-vi1-5X series false positive was the *uncultured human fecal virus* (UHFV, taxid 239364). UHFV was called with 15 contigs, which all originated from *Bifidobacterium pseudocatenulatum* assembly (GCF_022496265.1). We hypothesise that this may represent an unclassified bacteriophage genome. For the dog5X-ba.comp5X-vi5X metagenome false positive was the *Human gammaherpesvirus 4* (syn. *Epstein-Barr virus*, EBV) with just 18 reads. This originated from *Corynebacterium amycolatum* assembly (NZ_CP102778.1), which was added to RefSeq at a later time point (2022/08/23) than the compilation of our reference database (2022/06/20).

**Table 5.** Accessing *Lazypipe2 --ann minimap* accuracy on canine simulated metagenome. Here *ba5X* and *ba.comp5X* represent the canine bacterial genomes and canine complete bacterial genomes, respectively. True, number of correct taxa, TP, true positives, FP, false positives, FN, false negatives, Pr, precision, Rc, recall, F1, F1-metric.

| Metagenome | Target taxa | TRUE | TP | FP | FN | Pr | Rc | F1 |
| --- | --- | --- | --- | --- | --- | --- | --- | --- |
| dog5X-ba.comp5X-vi5X | Virus species | 35 | 35 | 1 | 0 | 97.2% | 100.0% | 98.6% |
| dog5X-ba5X-vi5X | Virus species | 35 | 35 | 1 | 0 | 97.2% | 100.0% | 98.6% |
| dog5X-ba5X-vi4X | Virus species | 35 | 35 | 1 | 0 | 97.2% | 100.0% | 98.6% |
| dog5X-ba5X-vi3X | Virus species | 35 | 35 | 1 | 0 | 97.2% | 100.0% | 98.6% |
| dog5X-ba5X-vi2X | Virus species | 35 | 35 | 1 | 0 | 97.2% | 100.0% | 98.6% |
| dog5X-ba5X-vi1X | Virus species | 35 | 34 | 1 | 1 | 97.1% | 97.1% | 97.1% |
| dog5X-ba.comp5X-vi5X | Bacterial species | 53 | 40 | 9 | 13 | 81.6% | 75.5% | 78.4% |
| dog5X-ba.comp5X-vi5X | Bacterial genera | 32 | 31 | 1 | 1 | 96.9% | 96.9% | 96.9% |
| dog5X-ba5X-vi5X | Bacterial species | 159 | 96 | 82 | 63 | 53.9% | 60.4% | 57.0% |
| dog5X-ba5X-vi5X | Bacterial genera | 71 | 61 | 3 | 10 | 95.3% | 85.9% | 90.4% |

Similar to benchmarking on human simulated metagenome, retrieval of bacterial taxa was evaluated ignoring predictions falling in the last 5 percentiles of read distribution. The accuracy for bacteria was comparable to human metagenome results. Recall and precision for bacterial genera were relatively high: 85.9-96.9% and 95.3-96.9%, respectively (Table 5).

### 3.2 Benchmarking read binning and genome coverage

We evaluated read binning and genome coverage from *Lazypipe2 --ann minimap* results for the canine simulated metagenome. For each viral genome we extracted ids for the simulated reads and reads assigned by the pipeline. From these numbers we estimated recall and precision for read taxonomic binning. For 64% of viral genomes both recall and precision for read binning exceeded 99% and for 85% of viral genomes these exceeded 80% (Table S1).

We also estimated horizontal coverage of viral genomes by the resulting assemblies. The pipeline created assemblies for 37 out of 39 unique virus taxids in the benchmark. The exceptions were the three canine parvovirus genomes that were all assembled to a single genome and reported as the *Canine parvovirus 2a*. Genome coverage for assembled viral genomes ranged between 58% and 100% with median at 92% (Figure 2).

**Figure 2.**
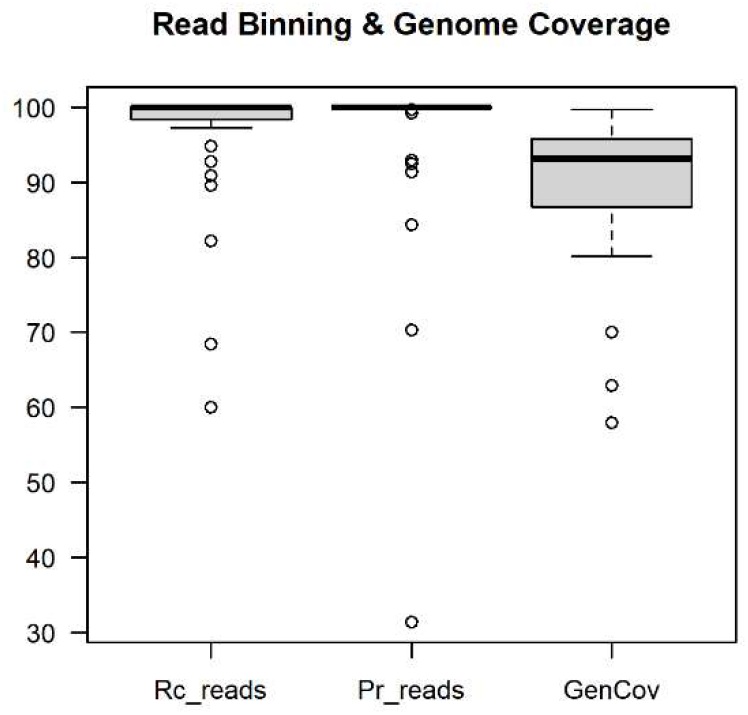
Benchmarking read binning and genome coverage. The first two boxplots represent recall and precision of read taxonomic binning for viral genomes. The last column represents coverage of viral genomes by the resulting assemblies.

### 3.3 Benchmarking time performance

We compared the execution time of Lazypipe2 (with options --*ann minimap* and --*ann sans*) to Lazypipe1 (with option –*ann sans*) and Kraken2. Benchmarking was done with human simulated metagenome [16] on Linux/Unix CPU supercluster with 32 cores each running at 2.1 GHz. Real and CPU times for the compared tools are displayed in Table 6. The new version of the pipeline was slightly slower than v1.0. This is mainly due to overhead introduced by fastq file compression. The main bottleneck was the database search. For both *Lazypipe1 –ann sans* and *Lazypipe2 –ann sans* database search accounted for 70-71% of real execution time.

**Table 6.**
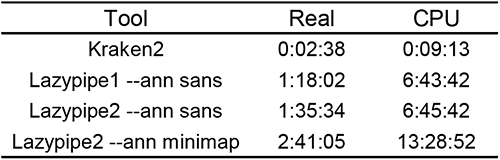
Benchmarking time performance. Real and CPU execution times for different tools with the human simulated metagenome.

## 4. DISCUSSION

Metagenomic analysis pipelines are vital for global pathogen detection and monitoring. Here, we presented Lazypipe2, an updated version of our mNGS pipeline with significant improvements in code ability, transparency and support for new software components. The previous version, Lazypipe, and now Lazypipe2 have been used contributing to virus discovery, and demonstrating its potential for unbiased NGS-based studies [22–26].

Benchmarking on simulated metagenomes demonstrated that assembling and taxonomic binning of contigs with minimap2 against a subset of NCBI nt is a highly accurate strategy for calling known viruses. For the human simulated metagenome Lazypipe2 --*ann minimap* achieved 94.0% recall and 97.5% precision for viral species. For the canine simulated metagenome Lazypipe2 --*ann minimap* had 100% recall and 97.2% precision. Notably, this high accuracy was sustained even with a heavy host and bacterial sequence background with viral reads spiked at just 1X coverage and constituting just 0.004% of the total NGS library. Additionally, we demonstrated that viral genomes assembled from viral reads spiked at 5X had good horizontal coverage (median at 92%). Also, recovery of the spiked reads for viral genomes was highly accurate (precision and recall exceeding 99%) for two thirds of viral genomes and at high level (>80%) for 85% of viral genomes spiked at 5X.

Annotating with SANSparallel (a homology search for aa sequences) had a slightly lower performance for calling known viruses from the human simulated metagenome. On this benchmark *Lazypipe2 --ann sans* showed 89.3% recall and 85.2% precision for viral species. For annotations based on aa sequences most errors were failures to identify measurable orfs and misalignments of ORFs to false positives. For nt-based annotations misalignment to false positives were less frequent. These observations support the choice of nt-based annotations for known viral targets with low divergence from reference sequences. Possible scenarios for applying nt-based annotations include surveillance of a list of known pathogens from various samples and diagnostics or research targeting known viruses with clinical samples.

Annotations based on aa sequences have higher sensitivity for viruses with higher divergence from the reference [2,17,40]. We also must consider that current reference databases are estimated to represent only a fraction of viral diversity [41]. These points advocate in favour of homology search with aa sequences when looking for novel and divergent viruses. However, there is a trade-off between finding potential new viruses with relatively low aa identity, and misclassification of host, environmental or bacterial sequences as potential viruses (false positives).

Identification of known viruses with *Lazypipe2 --ann minimap* approached perfect accuracy in detecting viruses from simulated metagenomes. The remaining errors were caused by filtering of retroviruses with the host reads and close homologs that were >99.9% identical within the assembled region. Correct identification of retroviruses prior to host filtering is an important goal for future development. This study also left out benchmarking on real datasets, although we expect the performance to be at least at the level of Lazypipe1 (tested on a mock-virome dataset).

Other important goals for future development is improving the detection of divergent novel viruses. This can be approached in several ways, for example, by integrating well established and highly sensitive techniques based on Hidden Markov Models for detecting protein homologs (e.g. HMMER [42]). Similar ideas have been implemented in other mNGS pipelines [40]. Higher sensitivity will pose challenges such as higher number of false positives and the nontrivial task of evaluating performance. Benchmarking the detection of novel divergent viruses is difficult to formulate, although some efforts have been made using methods for simulated evolution [2,40].

Most of the currently existing mNGS pipelines only supports short-read sequencing platforms but we are planning to work further on Lazypipe2 to support long-read platforms (e.g. Oxford Nanopore Technologies which is becoming highly popular for pathogen surveillance due to its portability and cost effectiveness) and make it more user friendly by developing a web interface for fairly self-explanatory results in Hypertext Markup Language. This will help scientists and clinicians with minimum bioinformatics skills to analyse their samples and gain insight from mNGS datasets for both known and novel pathogens.

## Author Contributions

Conceptualization of the project: R.K., T.S. and O.V.; Conceptualization of the pipeline: I.P.; Implementation, updates and code distribution: I.P.; Reference Database Curation: I.P.; Benchmarking: I.P..; Testing and Reporting: I.P., R.K. and T.S.; Writing – Original Draft Preparation: I.P., R.K. and T.S.; Writing – Review & Editing: O.V., R.K., T.S. and T.Si.; Supervision of the study: R.K., T.S. and O.V.; Funding Acquisition: O.V. and T.Si. All authors have read and agreed to the published version of the manuscript.

## Acknowledgements

The authors wish to acknowledge CSC – IT Center for Science, Finland, for computational resources.

## Data Availability Statement

Lazypipe2 user manual and reference databases are openly available at https://www.helsinki.fi/en/projects/lazypipe. Lazypipe2 source code is openly available from the git repository at https://bitbucket.org/plyusnin/lazypipe/.

## Supplementary Materials

File S1: Accession ids and other information for virus and bacterial genomes included in the canine simulated metagenome.

File S2: Misclassification errors by *Lazypipe2 --ann minimap* and --*ann sans* for human simulated metagenome.

Table S1: Results for viral read binning and genome coverage for canine simulated metagenome.

