## Supplementary material for "Enhanced Viral Metagenomics with Lazypipe 2": File S2

**File S2 for Enhanced Viral Metagenomics with Lazypipe2**

### Classification errors for human simulated metagenome

*Human endogenous retrovirus* and *Human endogenous retrovirus W* were not predicted. All reads from these genomes were filtered as host reads, since these aligned to the human genome.

*Naples phlebovirus* was misclassified as *Toscana phlebovirus* (false positive). All 280 reads labelled in the benchmark as *N. phlebovirus* were assembled to a single 4,187 nt contig (*k141.9952*) with a single 960 nt orf detected. This orf was mapped by the pipeline to *Toscana phlebovirus* with 99.3% coverage and 99.0% identity. Mapping with blastn confirmed 100% contig identity to *Toscana phlebovirus* M segment (acc X89628.1) and only with 82.1% identity to *Naples phlebovirus*. A blastn search with a sample of 28 reads also all mapped to the same *Toscana phlebovirus* M segment. In this case, sampled reads and assembled contig both had higher identity to the reported false positive.

*Mopeia Lassa virus reassortant 29* was misclassified as *Lassa virus* (false positive). All 220 reads were assembled to a single 3,375 nt contig (*k141.54025*) with a single 1,551 nt orf identified. This orf was mapped by the pipeline to *Lassa virus* (top 3 hits) with 99.6% coverage and 100% identity. Mapping with blastn confirmed 99.94% contig identity to *Lassa virus* and 99.88% identity to *Mopeia Lassa virus reassortant 29*. In this case, the sampled region also had higher identity to the false positive than the genome labelled as the ground truth.

*Hepatitis B virus* was a false negative missing from the result list. All 210 reads were assembled to a single contig of 3,173 nt (*k141.36746*) with a short 317 nt orf identified. The orf was erroneously mapped to *Trichinella spiralis.* Mapping with blastn confirmed 100% contig identity to *Hepatitis B virus*. The problem in this case was the lack of sufficiently longs orfs.

*Uukuniemi uukuvirus* was a false negative missing from the result list. All 210 reads were assembled to a single contig of 3,199 nt (*k141.29920*) with just two short orfs identified (132 nt and 279 nt). There were no significant database hits for these orfs. Mapping with blastn confirmed 100% contig identity to *Uukuniemi virus* (acc M17417.1).

*Influenza A virus* was a false negative missing from the result list. All 90 reads were assembled to a single contig of 1,386 nt (*k141.46069*) with a single 105 nt orf identified. There were no significant database hits for this orf. Mapping with blastn confirmed 100% identity to *Influenza A virus* (acc AJ404629.1).

*Mason-Pfizer monkey virus* (syn. *Human type D retrovirus*) was misclassified as *Squirrel monkey retrovirus* (false positive). All 65 reads were assembled to a single contig of 308 nt (*k141.29825*) with a single 306 nt orf identified. Pipeline mapped this orf to *Squirrel monkey retrovirus* (also a *Betaretrovirus*). However, this prediction was filtered from abundance estimation due to low read count. Mapping with blastn confirmed 100% identity to *Human type D retrovirus* (acc D10443.1).

Part of *Vaccinia virus* reads were misassigned to *Cowpox virus* (false positive). Cowpox was reported with 551 PE reads, and a single 174,612 nt contig (*k141.74634*) with 157 orfs. Pipeline mapped these orfs to various *Orthopox* viruses including *Camelpox virus*, *Cowpox virus, Monkeypox virus*, and *Vaccinia virus*. However, only *Cowpox* and *Vaccinia virus* were picked by the weighting model. Mapping with blastn confirmed 100% contig identity to *Vaccinia virus* (acc AY243312.1).

There was confusion with labelling *Borna disease viruses* (BDV). According to the original publication [1], the benchmark included 590 PE reads from *Borna disease virus* (taxid 12455). However, the corresponding reads were marked with *NC_001607* accession, which corresponds to *Borna disease virus 1* (taxid 1714621, considered here as the ground truth). Pipeline assembled 589 PE reads labelled with NC_001607 accession to a single contig (*k141.57245*) of 8,890 nt with three orfs. Three orfs mapped to *Borna disease virus* (taxid 12455, 98.6-99.5% query coverage and 100% identity) and two orfs mapped to *Borna disease virus 1* (taxid 1714621, 98.6-99.5% query coverage and 100% identity), which were both reported. Mapping with blastn confirmed 100% contig identity to *Borna disease virus 1* (taxid 1714621).

Part of *Simian foamy virus (SFV)* reads were misclassified to *Central cimpanzee SFV* (false positive). This was reported with 193 PE reads, one contig (*k141.72284*) and two orfs. These orfs were mapped by the pipeline to *SFV* (99.6% coverage and 99.0-100% identity) and *Central cimpanzee SFV* (99.1-99.6% coverage and 96.0-99.0% identity). Mapping with blastn confirmed 99.00-99.15% contig identity to *Human spumaretrovirus*, 98.50% identity to *Central cimpanzee SFV* (false positive) and 96.71%-98.21% identity to *SFV* (ground truth).

Part of *SVF* reads were also misclassified to *African green monkey SVF* (false positive). This was reported with 32 PE reads, one contig and one orf 753 nt. This orfs was mapped to *African green monkey SVF* with 97.2% coverage and 81.0% identity.

Another false positive was the *Eastern chimpanzee SFV*. This was reported with 146 PE reads, three contigs (*k141.47714*, *k141.38283* and *k141.9270*) and four orfs. Mapping with blastn showed 99%-100% contig identity to *Human spumavirus* (not in the benchmark), 98.41-99.61% identity to the *Eastern chimpanzee SFV* and 85.03-96.75% identity to *SFV*.

Part of *Primate bocavirus 1* reads were misclassified to *Human bocavirus* (false positive) and *Bocaparvovirus sp.* (false positive, taxid 1883111). These were reported with 154 and 36 PE reads, one 5,252 nt contig (*k141.64308*) and one to three orfs. Pipeline mapped three orfs to *Primate bocavirus 1*, three orfs (1,920 nt, 441 nt and 2,016 nt) to *Human bocavirus* (with 100% identity) and one orf (1,920 nt) to *Bocaparvovirus sp*. (100% identity). Mapping with blastn confirmed 100% contig coverage and identity to the ground truth *Primate bocaparvovirus 1*, 99.94% identity to *Human bocavirus* (false positive) and 99.90% identity to *Bocaparvovirus sp*. (false positive).

*Chimeric Tick-borne encephalitis virus/Dengue virus* 4 (false positive, taxid 638787) was reported with 349 PE reads, one 10,624 nt contig (*k141.780*) and a single 6,108 nt orf. Pipeline mapped this orf to *Dengue virus 4* and the chimeric virus with identical scores, 99.9% query coverage and 100 % identity. Mapping with blastn showed that the contig was 100% identical to a fragment of *Dengue virus 4* genome. The error here is due to the orf beeing identical between the two viruses.

*Phlebovirus SDYY104/China/2011* virus (false positive, taxid 1848960) was reported with 140 PE reads, one 6,353 nt contig (*k141.30190*) and a single 6,258 nt orf. Pipeline mapped this orf to *Phlebovirus SDYY104/China/2011* with 99.9% coverage and 100% identity. Mapping with blastn confirmed 100% identity of the contig to *Dabie bandavirus*.

Part of *Dabie bandavirus* reads were misclassifed to *SFTS phlebovirus* (false positive, taxid 1933190). This was reported with 88 PE reads, one 1,700 nt contig (*h*), and a single 738 nt orf. Pipeline mapped this orf to *SFTS phlebovirus* with 99.2% coverage and 100% identity.
