## Supplementary material for "Enhanced Viral Metagenomics with Lazypipe 2": Table S1

| Virus | Taxid | Spike dcov | Spike readn | Rc | Pr | GenCov |
|---|---|---|---|---|---|---|
| Astrovirus dogfaeces/Italy/2005 | 566308 | 5X | 45 | 82.2% | 92.5% | 80.1% |
| Betapolyomavirus canis | 1980633 | 5X | 83 | 100.0% | 100.0% | 89.4% |
| Canid alphaherpesvirus 1 (CHV) | 170325 | 5X | 2085 | 99.9% | 100.0% | 86.7% |
| Canine adenovirus 1 | 10512 | 5X | 1016 | 99.9% | 99.7% | 99.7% |
| Canine adenovirus 2 | 10514 | 5X | 520 | 99.4% | 100.0% | 97.0% |
| Canine astrovirus | 1157338 | 5X | 110 | 97.3% | 93.0% | 89.1% |
| Canine bocavirus 1 | 1511885 | 5X | 90 | 100.0% | 100.0% | 93.7% |
| Canine bocavirus 3 | 1295080 | 5X | 88 | 100.0% | 100.0% | 93.7% |
| Canine circovirus | 1194757 | 5X | 33 | 90.9% | 100.0% | 70.0% |
| Canine kobuvirus | 1836608 | 5X | 135 | 89.6% | 70.3% | 82.9% |
| Canine kobuvirus CH-1 | 1281440 | 5X | 135 | 60.0% | 84.4% | 58.0% |
| Canine kobuvirus US-PC0082 | 1082189 | 5X | 138 | 92.8% | 91.4% | 84.4% |
| Canine minute virus | 329639 | 5X | 83 | 100.0% | 100.0% | 95.9% |
| Canine morbillivirus | 11232 | 5X | 260 | 100.0% | 100.0% | 96.9% |
| Canine parvovirus | 10788 | 5X | 88 | 0.0% | 0.0% | 0.0% |
| Canine parvovirus 2a | 497961 | 5X | 83 | 100.0% | 31.4% | 95.8% |
| Canine parvovirus 2b | 568986 | 5X | 83 | 0.0% | 0.0% | 0.0% |
| Canine picodicistrovirus | 1150861 | 5X | 145 | 100.0% | 100.0% | 95.9% |
| Canine picornavirus | 1196647 | 5X | 130 | 100.0% | 100.0% | 92.3% |
| Canine respiratory coronavirus | 215681 | 5X | 515 | 94.8% | 100.0% | 87.7% |
| Canine vesivirus | 1176484 | 5X | 140 | 97.9% | 100.0% | 85.6% |
| Canis familiaris papillomavirus 13 | 1226723 | 5X | 135 | 98.5% | 100.0% | 89.2% |
| Canis familiaris papillomavirus 2 | 2759772 | 5X | 135 | 100.0% | 100.0% | 93.5% |
| Canis familiaris papillomavirus 3 | 360397 | 5X | 130 | 100.0% | 99.2% | 94.6% |
| Canis familiaris papillomavirus 4 | 464980 | 5X | 128 | 100.0% | 100.0% | 94.3% |
| Canis familiaris papillomavirus 6 | 1513269 | 5X | 135 | 100.0% | 100.0% | 96.3% |
| Canis familiaris papillomavirus 8 | 1081055 | 5X | 128 | 98.4% | 100.0% | 90.0% |
| Deltapolyomavirus canis | 2170403 | 5X | 85 | 100.0% | 100.0% | 96.3% |
| Faeces associated gemycircularvirus 1 | 1843735 | 5X | 35 | 100.0% | 100.0% | 89.3% |
| Influenza A virus (H3N2) | 11320 | 5X | 213 | 100.0% | 100.0% | 88.4% |
| Lambdapapillomavirus 2 | 35258 | 5X | 143 | 99.3% | 100.0% | 94.6% |
| Lupine bocavirus | 2017714 | 5X | 85 | 100.0% | 100.0% | 97.1% |
| Lyssavirus rabies | 11292 | 5X | 198 | 99.5% | 100.0% | 93.2% |
| Norovirus dog/GVI.1/HKU_Ca026F/2( | 673457 | 5X | 125 | 100.0% | 100.0% | 94.6% |
| Parainfluenza virus 5 | 2905673 | 5X | 253 | 68.4% | 100.0% | 62.9% |
| Picobirnavirus dog/KNA/2015 | 1961162 | 5X | 28 | 100.0% | 100.0% | 86.0% |
| Pneumovirus dog/Bari/100-12/ITA/2( | 2482952 | 5X | 248 | 100.0% | 100.0% | 97.6% |
| Rotavirus I | 1637496 | 5X | 288 | 99.7% | 100.0% | 86.1% |
| Torque teno canis virus | 687385 | 5X | 45 | 100.0% | 100.0% | 93.8% |

**Table S1.** Benchmarking read binning and genome coverage. Viral genomes reported by Lazypipe2 *–ann minimap* for the canine simulated metagenome. Spike dcov, depth of genome coverage set in ARK simulation, Spike readn, number of reads output by ARC simulation, Rc, recall, Pr, precision, GenCov, genome coverage by the assembly.
